# Evaluation of Three Forms of *Rhizoctonia solani* Mediated Pathogenicity to Sugar Beet Cultivars in Greenhouse Studies

**DOI:** 10.1101/2022.02.20.481202

**Authors:** Md Ehsanul Haque, Most Shanaj Parvin

## Abstract

*Rhizoctonia solani* causes damping-off, as well as crown and root rot of sugar beet (*Beta vulgaris* L). This pathogen overwinters as sclerotia or melanized mycelia. Traditionally, the resistance of cultivars to *R. solani* is evaluated by scoring disease reactions of the crowns and roots of older seedlings, instead of evaluating at seed germination. Most studies that have evaluated cultivar resistance to *R. solani* have used colonized whole barley grains as artificial inocula. Colonized grains are prone to contamination with other pathogens and are often lost to rodents/birds when applied in the field. Considering those limitations, a study was undertaken (1) to develop in vitro methods to generate natural sclerotia of *R. solani* on a large scale, (2) to compare pathogenic potentials of *R. solani* sclerotia, mycelia, and colonized barley grains for optimization of damping-off assays, and (3) to evaluate resistance of selected commercial cultivars to *R. solani*. Of the six-culture media, amended clarified V8 (ACV8) was the most suitable culture medium to grow and produce sclerotia on a large scale and 10% PDA was the least suitable. Three testing sizes of sclerotia were found to be equally effective in causing plant losses. Sclerotia inocula were comparable with mycelial discs and colonized barley grains in causing pre-emergence damping off under aseptic in vitro conditions. Sclerotia also were equally or more effective than mycelia plug or barley grain inocula in reducing seedling emergence, inducing damping off, and increased root rot under greenhouse in vivo conditions. To conclude, sclerotia can be prepared on a feasible scale and used as natural inocula to screen response to *R. solani* on sugar beet.

## 1. Introduction

Sugar beet (*Beta vulgaris*, L.) contributes approximately 20% of worldwide sugar production while sugarcane contributes the rest (Dohm et al. 2014). In the US, sugar beet accounts for 55% of total sugar production (USDA, ERS, 2019). *Rhizoctonia solani* Kühn is among the soil-borne pathogens that affect sugar beet stands and sugar yields. This genetically complex soil-borne fungus causes pre- and post-emergence damping-off, and root and crown rot (Windels and Nabben 1989; Harveson et al., 2006). *R. solani* has 13 anastomosis groups (AGs; AG 1 to AG 13), some of which are host-specific while others have wide host ranges (Carling et al. 2002; Parmeter et al. 1969). Sugar beet is prone to the AG 2-2 strain (Parmeter et al. 1969). The main AG subgroup that causes significant yield losses is AG-2-2 IIIB (Brantner and Windels 2009; Windels et al. 1997). The primary inocula of the pathogen in nature are mycelia and sclerotia. This pathogen can survive in the soil for many years in the form of sclerotia (Sherwood 1967; Adams and Papavizas 1970; Papavizas 1970). The sclerotia germinate under humid conditions and are often stimulated to germinate by the root exudates of seedlings (Flentje et al. 1963). The host-pathogen interactions are generally initiated by mycelia that penetrate into the root cortex causing infections to the tissue (Armentrout and Downer 1987; Christou 1962). Rhizoctonia symptom phenotyping in the field often varies due to biotic and abiotic factors (Bolton et al. 2010; Behn et al. 2012). As a result, greenhouse evaluations using colonized-barley-grains are usually performed as a complement to field evaluations (Ruppel et al. 1979; Behn et al. 2012). Colonized-barley-grain preparation requires time and space, technical know-how to produce on a large scale, as well as protection from other air-borne pathogens (Paulitz 2002; Paulitz and Schroeder 2005; Mahmoudi and Ghashghaie 2013; Webb and Calderon 2015). It can also be difficult to relate disease severities obtained with colonized-barley-grains inocula in greenhouse studies to those results with colonized-barley-grains inocula in evaluation of sugar beet cultivars in the fields. Since *R. solani* rarely produces any basidiospores, it is problematic to use the same amount and/or concentration of fungal biomass on barley seeds for each evaluation (Cubeta and Vilgalys 1997). Using quantifiable sclerotia as inocula may circumvent those limitations. This study evaluated the morphology and development of sclerotia on six different artificial media and then compared Rhizoctonia sclerotia, mycelia and colonized barley grains for infection severity on sugar beet cultivars in the greenhouse.

## 2. Materials and Methods

### 2.1 Fungal Isolates of *R. solani*

Five *R. solani* isolates were collected from affected sugar beet tap roots from a field in 2018 in Hickson, North Dakota (ND), USA. Genomic DNAs (Norgen Biotek Corp, Fungi DNA Isolation Kit #26200) (Table B.1.) of the five isolates were used for polymerase chain reaction (PCR) with the internal transcribed spacer (ITS) (Sharon et al., 2008). Subsequently, PCR products were flushed by E.Z.N.A ®Cycle Pure Kit Omega Bio-tek (Norcross, GA) and four samples were sequenced by GenScript (Piscataway, NJ). The sequences were identical, and BLASTn analysis showed 100% sequence homology to *R. solani* AG 2-2 IIIB (Genbank accession: MN128569). Those isolates were maintained on an amended clarified V8 (ACV8) culture medium.

### 2.2 Evaluation of Culture Media for Production of Sclerotia of *R. solani*

Microbial media play a significant role in the optimum mycelial growth and differentiation of different fungal species. Six media: amended clarified V8 (ACV8), 50% potato dextrose agar (50% PDA), 10% PDA, methylene-benomyl-vancomycin (MBV), cornmeal agar (CMA) and water agar (WA), were prepared following the “Manual of Microbiological Culture Media” (Difco and BBL Manual 2009) (Table B. 2.). WA media used as control as it does not produce any sclerotia. The experimental design was a completely randomized design (CRD) with four replications. Mycelial discs (4.5 mm, Cork Borer) of *R. solani*, AG 2-2IIIB cut from the 7-day old mother colony (50% PDA) were transferred onto each of the six media (Table B. 3). After inoculation and plates were sealed with parafilm, the plates were incubated at 23±2 ºC in incubator. This experiment was conducted twice. Radial growth (mm) was measured using a digital caliper (Pittsburg 6” Composite DC, Item 93293) at four time points after transfer: 2-day, 4-day, 6-day, and 8-day, respectively. The number of sclerotia was counted at four-time points after transfer: 7-day, 14-day, 21-day, and 28-day, respectively.

### 2.3 Evaluation of Sclerotia Size Effects on Damping-off

Three groups of sclerotia were categorized with a measuring scale based on their size: large (≥4.00 mm), medium (<4.00 mm but ≥2.00 mm), and small (<2.00 mm but ≥0.5 mm). To evaluate the effect of sclerotia size on inoculum potential, an experiment was conducted in a humidity chamber at 25°±2º C and 85% relative humidity. Plastic pots (27 × 13 × 13 cm, T.O. Plastics, Inc.; Clearwater, MN, USA) were filled with vermiculite and perlite mixer (PRO-MIX FLX) amended with osmocote (N-P-K:15-9-12) fertilizer (Scotts Company; Marysville, OH). Ten sugar beet seeds (Crystal 101RR) were sowed at 2 cm deep furrow at 1 cm apart (Noor and Khan 2015) for each treatment. One sclerotium was placed, next to each seed at the same depth and covered with mixer. Three-size groups of sclerotia were used exclusively in furrow to inoculate sugar beet cultivar Crystal 101RR (susceptible check), with a completely randomized design of four treatments (including non-inoculated check) with four replications. Four plastic pots were considered for each treatment/category. This experiment was conducted twice. The seedlings damping-off were counted at 14 days post-inoculation (dpi).

### 2.4 In vitro Inoculation on PDA Using Three Forms of *R. solani* Inocula

To compare the efficacies of the three forms of inocula-sclerotia, mycelia and colonized-barley grains-sugar beet seeds of Crystal 101RR were co-cultured with each form of inoculum on PDA with four replications. A non-inoculated check with four replications was used as a control. Sugar beet seeds were washed with 70% ethanol for 1 minute and rinsed twice with sterile water. Seeds were dried on sterile blotter paper under the laminar airflow cabinet. Three seeds were placed with sterile forceps at 1 cm apart on each culture plate followed by each form of inocula being placed close to each seed. This experiment was conducted in a growth chamber at 25°±2º C. This experiment was conducted twice. Germination data were recorded at 7 days post inoculation (dpi).

### 2.5 Greenhouse Evaluation of Cultivars’ Susceptibility to *Rhizoctonia* Inocula

To determine if inoculum type had an effect on cultivar response to Rhizoctonia root rot, seven commercial sugar beet cultivars and four forms of inocula were arranged in CRD with four replications in a greenhouse. Root rot ratings (figures in brackets) of these seven commercial cultivars were reported (sbreb.org/research/, Research Report 2018) as follows: Crystal 101RR (4.50), Crystal 467RR (3.94), Hilleshog 4302RR (3.71), Maribo MA 504 (4.25), BTS 8606 (4.24), BTS8500 (4.36) and BTS80RR52 (3.96). Three forms of *Rhizoctonia* inocula, colonized-barley grains, sclerotia, and mycelia plug, along with autoclaved-barley grains as a control, were used to inoculate each cultivar. Plastic pots (27 × 13 × 13 cm, T.O. Plastics, Inc.; Clearwater, MN, USA) were filled with vermiculite and perlite mixer (PRO-MIX FLX) amended with osmocote (N-P-K:15-9-12) fertilizer (Scotts Company; Marysville, OH). Ten sugar beet seeds of each cultivars were sowed at 2 cm deep furrow at 1 cm apart (Noor and Khan 2015). One colonized-barley grain, one sclerotia, one mycelial plug and one autoclaved barley seed as mock inoculation was placed, respectively, next to each seed at the same depth and covered with mixer. The greenhouse temperature during the experiment period was 25 ± 2ºC, with 80% relative humidity, and a 12-hour photoperiod. Plants were watered as needed to maintain adequate soil moisture conducive for plant growth and disease development.

The seedling emergence, and damping-off were recorded at 14 days post-inoculation (dpi) and 42 dpi, respectively. At 56 dpi, plants were removed from pots, and roots were washed and rated for root rot disease severity using a modified 1-7 rating scale, where 1 = clean roots and no infection, 2 ≤ 5% of root surface with black/brown symptoms, 3 = 5-25% of root surface with black/brown symptoms; similarly, 4 = 26-50%, 5 = 51-75%, 6 = 75-100% of root surface with black/brown symptoms, and 7 = dead plants (withered) (Ruppel et al., 1979).

### 2.6 Statistical Analyses

Experiments were conducted twice as a complete randomized design (CRD) with four replicates. Levene’s test of homogeneity of variances was performed across the experiments before the data were combined. Data in all the experiments were analyzed using R-studio (Version 3.6.1, St. Louis, Missouri, USA). The data were subjected to analysis of variance (ANOVA) and Fisher’s Protected Least Significant Difference (LSD) was used to separate treatment means using the same R-package (3.6.1). Treatment means were distinguished by the calculated Fisher’s LSD at p = 0.05 probability level. One-way non-parametric ANOVA (Kruskal-Wallis test and Pairwise. Wilcox. Test) in R was performed for root rot ratings across forms of inocula and cultivars.

## 3. Results

### 3.1 Culture Media Suitability for Radial Growth and Sclerotia Production

Simple linear regressions of mean radial growth were calculated on days from transferring mycelial discs unto each of the six test culture media. Mean numbers of sclerotia were first analyzed using natural logarithmic transformations and then simple linear regressions of these logarithmically transformed sclerotia numbers were calculated. To test the difference in effects of culture media, comparisons of line positions of the intercept and/or line parallelisms of the slope were carried out based on a t-test. The separately fitted simple linear regression lines explained 96.6% of the variance in measured radial growth (i.e. R2=0.966, df=12) and 99.6% of the variance in natural logarithms of sclerotia numbers (i.e. R2=0.996, df=12). The estimated parameter values of intercept and slope in the fitted simple linear regression lines for radial growth and natural logarithms of sclerotia numbers are given in Table 1.

**Table 1.** Parameter estimates of intercepts and slopes in fitted simple linear regression lines for radial growth (mm) and natural logarithm of sclerotia numbers in six growing media.

| Indicator | Media<sup>a</sup> | Intercept | Slope |
| --- | --- | --- | --- |
| Radial growth<sup>b</sup> | ACV8 | 2.300 | 0.850 |
|  | 50% PDA | 0.450 | 1.020 |
|  | 10% PDA | 0.350 | 0.950 |
|  | MBV | -0.550 | 1.105 |
|  | CMA | 0.500 | 0.940 |
|  | WA | 0.050 | 0.925 |
| Sclerotia number<sup>c</sup> | ACV8 | 2.569 | 0.096 |
|  | 50% PDA | 1.772 | 0.095 |
|  | 10% PDA | 1.591 | 0.052 |
|  | MBV | 2.403 | 0.056 |
|  | CMA | 2.227 | 0.093 |
|  | WA | 0.000 | 0.000 |

<sup>a</sup>“ACV8” is amended clarified V8, “50% PDA” is 50% potato dextrose agar, “10% PDA” is 10% potato dextrose agar, “MBV” is methylene-benomyol-vancomycin, “CMA” is cornmeal agar and “WA” is water agar.
<sup>b</sup>Comparisons of line positions of the intercept and line parallelisms of the slope showed that ACV8 was significantly better than 10% PDA, 50% PDA, MBV, CMA and WA ( $p < 0.05$ ) in favor of mycelial growth. However, the radial growth was not significantly different ( $p > 0.05$ ) between any comparison pairs of 10% PDA, 50% PDA, MBV, CMA and WA.
<sup>c</sup>Comparisons of line positions of the intercept and line parallelisms of the slope showed that ACV8 was significantly better than 10% PDA, 50% PDA, MBV and WA ( $p < 0.05$ ), but better than CMA only at $p < 0.07$ . The number of sclerotia was significantly different between any comparison pairs of 10% PDA, 50% PDA, MBV, CMA and WA ( $p < 0.05$ ).

Results showed that ACV8 was significantly better than 10% PDA, 50% PDA, MBV, CMA and WA (p<0.05) in favor of mycelial growth (Table 1, Fig. 1). However, the radial growth was not significantly different between all comparison pairs of 10% PDA, 50% PDA, MBV, CMA and WA (p>0.05). For production of sclerotia numbers, ACV8 was significantly better than 10% PDA, 50% PDA, MBV and WA (p<0.05), but better than CMA only at p<0.07. The number of sclerotia was significantly different between all comparison pairs of 10% PDA, 50% PDA, MBV, CMA and WA (p<0.05). The visual differences in formation of sclerotia were shown in Fig. 2 on six culture media at 28 days after transfer.

**Fig. 1.**
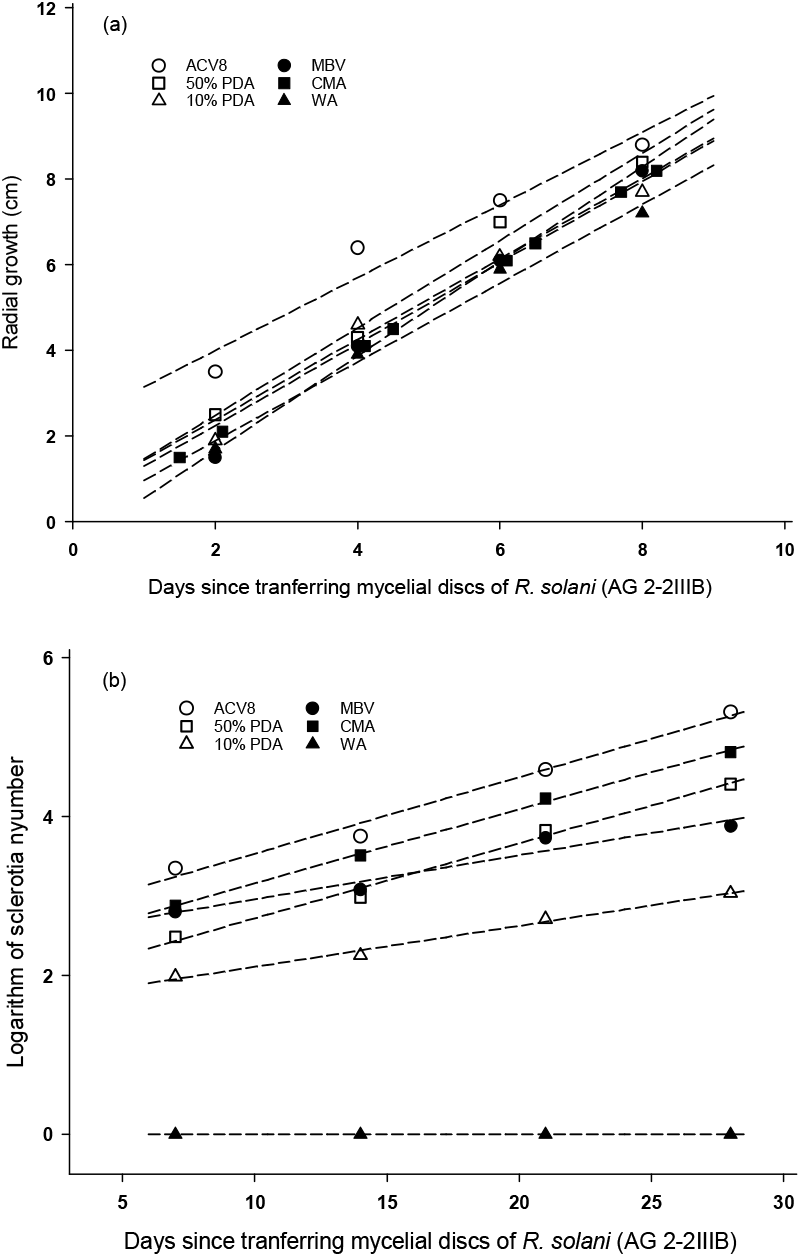
Increases in radial growth (a) and natural logarithm of sclerotia numbers (b) after days from transferring mycelial discs of *R. solani* (AG 2-2IIIB) unto six culture media. Broken lines are fitted simple linear regressions with intercepts and slopes and their tests are shown in Table 1. “ACV8” is amended clarified V8, “50% PDA” is 50% potato dextrose agar, “10% PDA” is 10% potato dextrose agar, “MBV” is methylene-benomyl-vancomycin, “CMA” is cornmeal agar and “WA” is water agar.

**Fig. 2.**
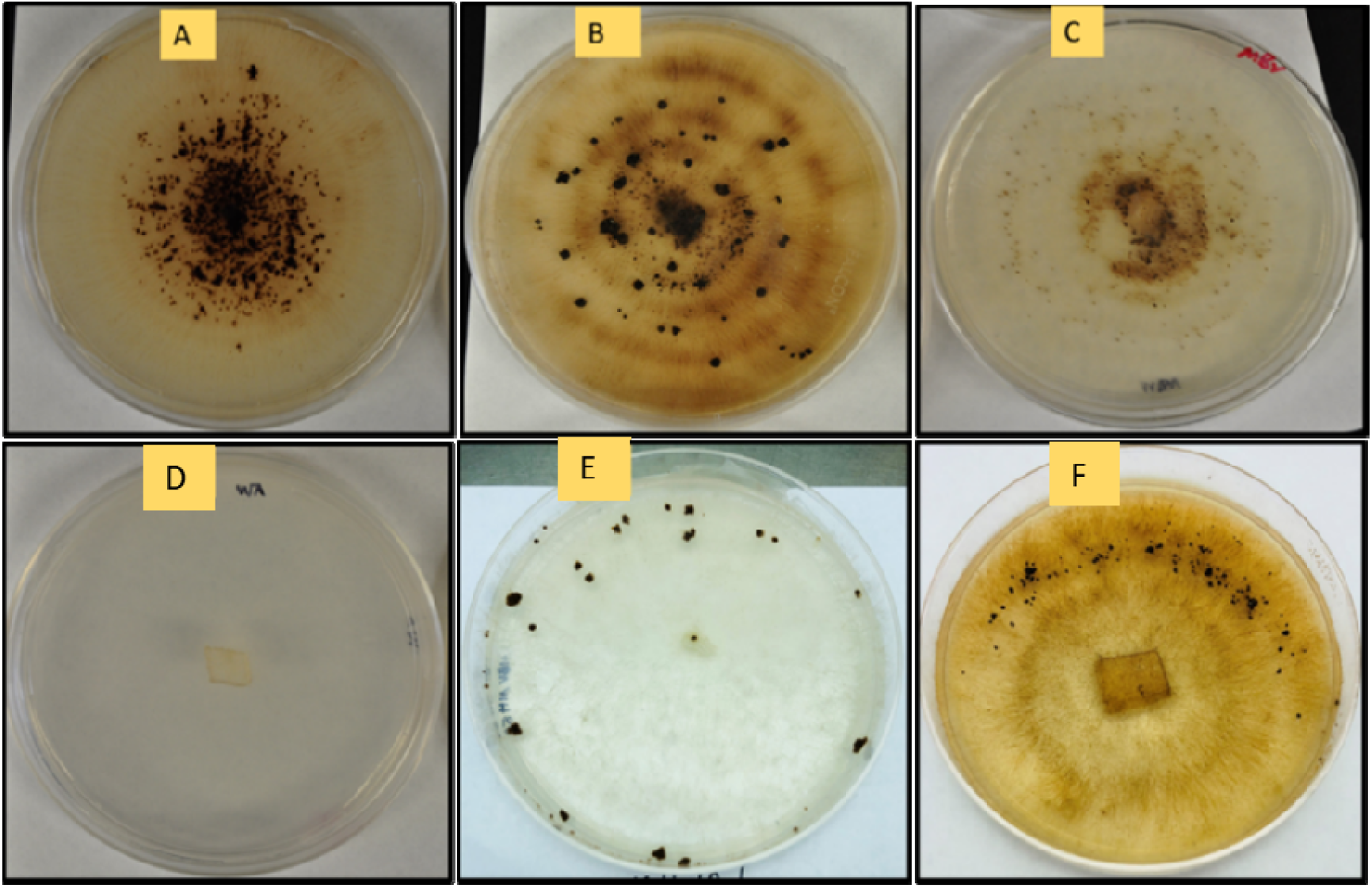
Visual difference in development and formation of sclerotia on six culture media at 28 days after transfer. Cultural media are: A: amended clarified V8 (ACV8), B: CMA: cornmeal agar (CMA), C: MBV: methylene-benomyl-vancomycin (MBV), D: WA: water agar (WA), E: 10% PDA: 10% potato dextrose (10% PDA), and F: 50% PDA: 50% potato dextrose agar (50% PDA).

### 3.2 Inoculum Potentials of Different Sizes of Sclerotia

Mean comparison tests showed that all sizes of sclerotia significantly reduced plant stands at 14 dpi compared with the control treatment (p<0.05), and they all caused the same 60% of plant losses (p>0.05) (LSD = 1.15 at p=0.05) (Fig. 3). The inoculum potentials of sclerotia sizes had equal capacities to infect sugar beet seedlings.

**Fig. 3.**
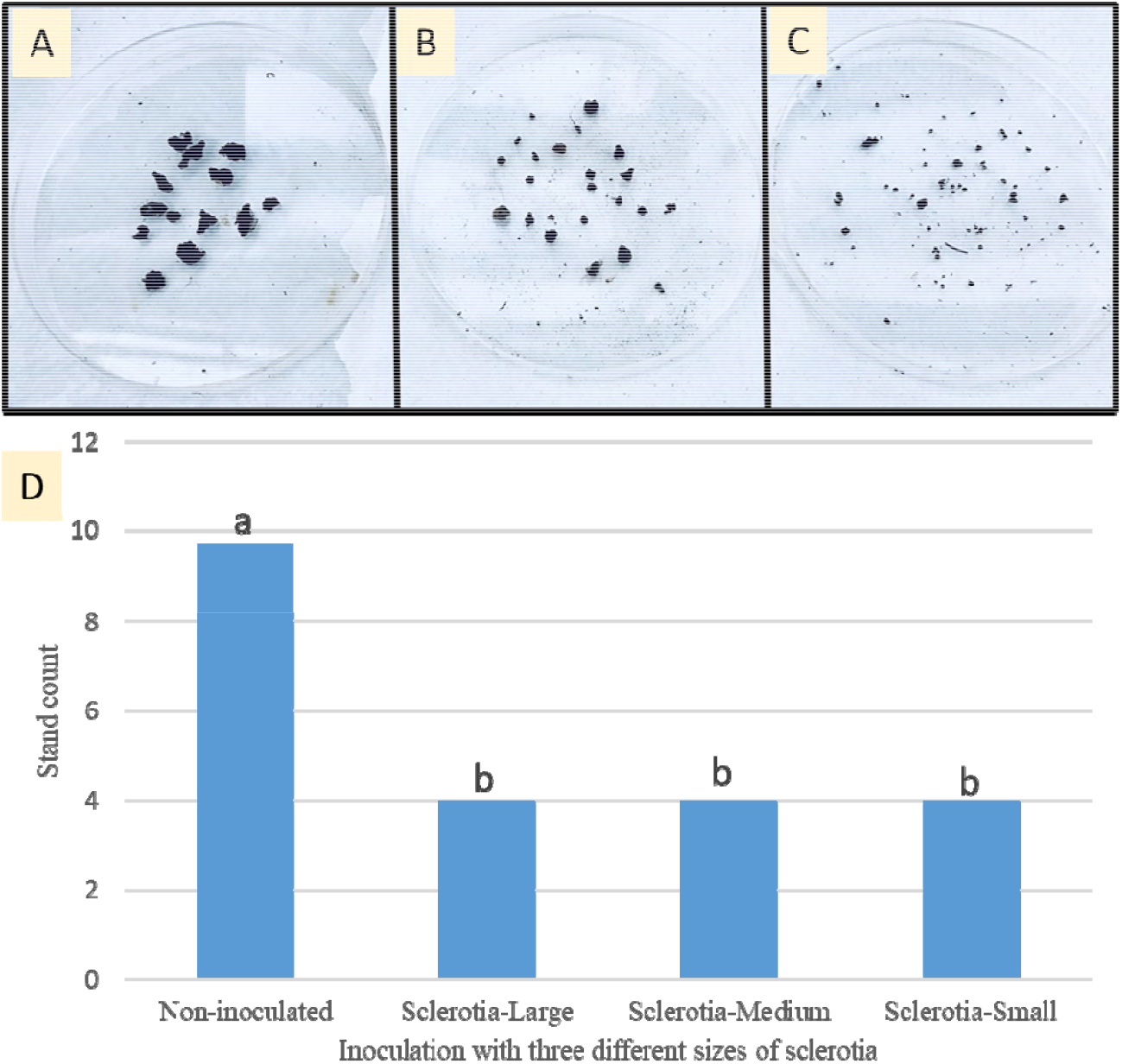
Three different categories of sclerotia and its pathogenicity test. A, Large (≥4.0 mm). B, (<4.00 mm but ≥2.00 mm). C, small (<2.00 mm but ≥0.5 mm) and D, stand count at 2 weeks of post-inoculation. Mean followed by the same letters are not significantly different at α = 0.05 at which LSD 1.15, and MSE =0.562.

### 3.3 Observing Damping-off In vitro

The three different forms of *R. solani* inocula tested all resulted in visual damping-off at 7 dpi, while 100% seedling emergence was observed in the non-inoculated check (Fig. 4). All three inoculum types yielded a similar severity of damping-off symptoms under in vitro conditions.

**Fig. 4.**
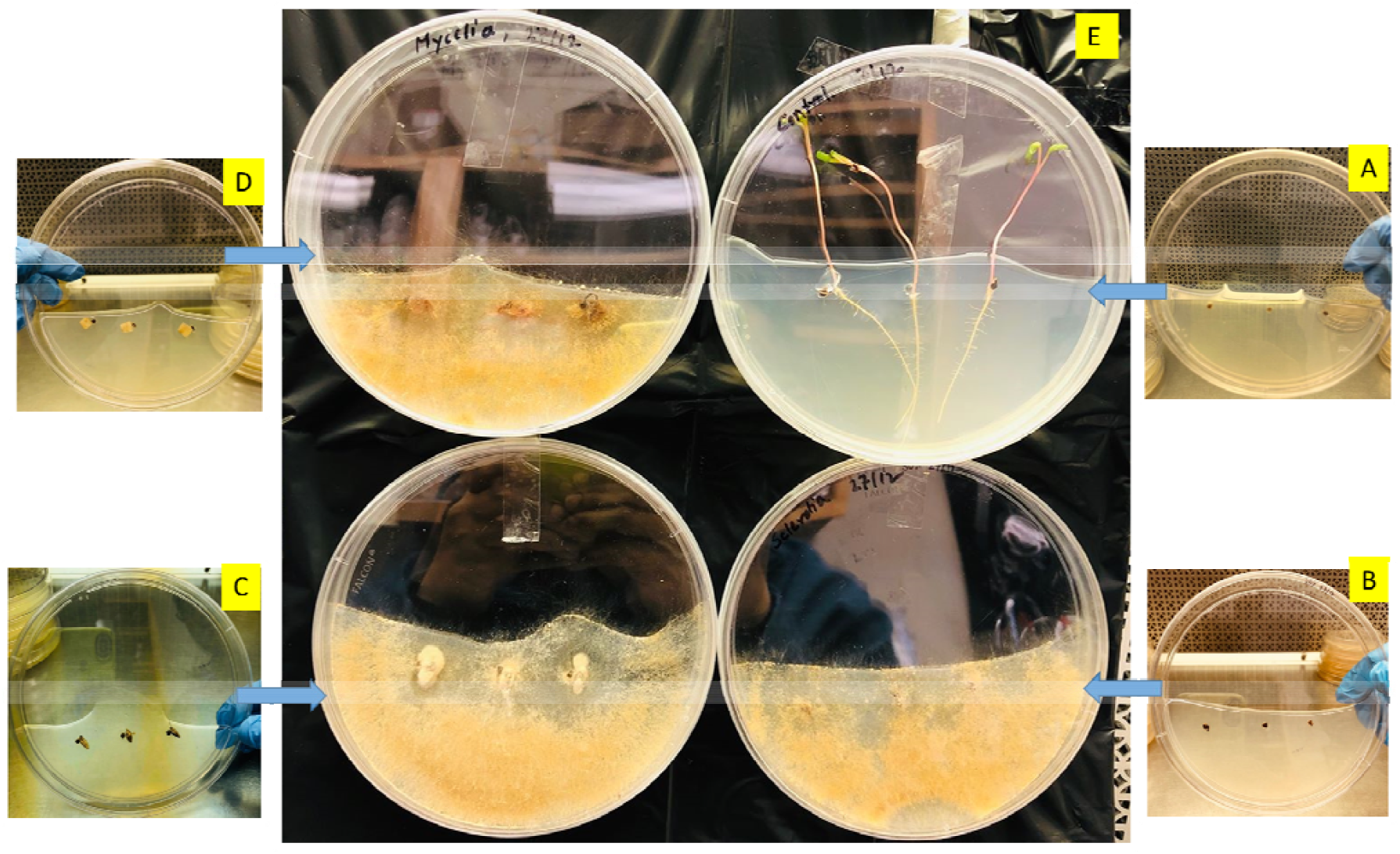
In vitro inoculation on PDA media using three forms of *R. solani* inocula and a control: A: with only seed (control check), B: with seed and sclerotia, C: with seed and colonized barley grains, and D: with seed and mycelia plug, E: Center picture illustrates the corresponding pre-emergence damping-off at 7 dpi. Only non-inoculated (control) resulted in seedling development.

### 3.4 Effect of Forms of *Rhizoctonia* Inoculum on Sugar beet Cultivar Response in the Greenhouse

The effects of inoculum forms on cultivars were significantly different on percentage of emergence at 14 dpi (p<0.05) and percentage damping off at 42 dpi (p<0.05) (Table 2). Among the three inoculum forms, sclerotia inoculum resulted in the lowest mean emergence (42.86%)) when averaged across all cultivars, with the non-inoculated check showing the highest mean emergence (96.07%) Colonized barley grains and mycelial forms resulted overall emergence 64.29% and 54.29%, respectively for all cultivars (Table 2). The highest mean emergence was observed in Maribo MA 504 (72.50%), followed by BTS 8500 (69.38%). There was overall similar significance level between in the mean emergence of BTS 80RR52 and BTS 8600 (66.25%). The lowest mean emergence was found in Crystal 101RR (54.38), which was followed by Crystal 467RR (58.13%).

**Table 2.** Percentage of seedling emergence at 14 dpi and percentage of damping-off at 42 dpi. Means followed by the same letters are not significantly different at p=0.05.

| Indicator | Cultivar | Forms of inoculum |  |  |  | Cultivar mean |
| --- | --- | --- | --- | --- | --- | --- |
|  |  | Colonized barley grains | Mycelia | Sclerotia | Autoclaved barley grains |  |
| Emergence (%) | BTS80RR52 | 62.50 | 55.00 | 52.50 | 95.00 | 66.25b |
|  | BTS8500 | 62.50 | 62.50 | 55.00 | 97.50 | 69.38ab |
|  | BTS 8606 | 65.00 | 57.50 | 45.00 | 97.50 | 66.25b |
|  | Crystal 101RR | 55.00 | 45.00 | 22.50 | 95.00 | 54.38d |
|  | Crystal 467RR | 65.00 | 45.00 | 27.50 | 95.00 | 58.13cd |
|  | Hilleshog 4302RR | 65.00 | 55.00 | 37.50 | 97.50 | 63.75bc |
|  | Maribo MA 504 | 75.00 | 60.00 | 60.00 | 95.00 | 72.50a |
|  | **Inoculum mean** | 64.29b | 54.29c | 42.86d | 96.07a |  |
| Damping-off (%) | BTS80RR52 | 52.5 | 57.5 | 67.5 | 0.0 | 44.38d |
|  | BTS8500 | 72.5 | 72.5 | 72.5 | 0.0 | 54.38a |
|  | BTS 8606 | 57.5 | 67.5 | 80.0 | 0.0 | 51.25b |
|  | Crystal 101RR | 60.0 | 75.0 | 85.0 | 0.0 | 55.01a |
|  | Crystal 467RR | 50.0 | 65.0 | 75.0 | 0.0 | 47.50bc |
|  | Hilleshog 4302RR | 47.5 | 77.5 | 67.5 | 0.0 | 48.12bc |
|  | Maribo MA 504 | 40.0 | 65.0 | 72.5 | 0.0 | 44.37d |
|  | **Inoculum mean** | 54.29c | 68.58b | 74.29a | 0.0d |  |

**Table 3.** Non-parametric analysis for root rot ratings* at 56 dpi caused by forms of inocula

| Inocula | Mean Rank Severity | Variance | Lower Limit | Upper Limit | Min | Max |
| --- | --- | --- | --- | --- | --- | --- |
| Autoclaved barley grains | 1 | 0.0 | 0.64 | 1.35 | 1 | 1 |
| Colonized barley grains | 2.31 | 0.22 | 1.97 | 2.71 | 2 | 3 |
| Mycelia | 3.07 | 0.50 | 2.81 | 3.41 | 2 | 5 |
| Sclerotia | 3.75 | 0.50 | 3.35 | 4.07 | 2 | 7 |

\*Ratings were 1 = clean roots and no infection, 2 ≤ 5% of root surface with black/brown symptoms, 3 = 5-25% of root surface with black/brown symptoms; similarly, 4 = 26-50%, 5 = 51-75%, 6 = 75-100% of root surface with black/brown symptoms, and 7 = dead plants (withered) (Ruppel et al. 1979).

Among the three forms of *Rhizoctonia* inocula, the highest damping-off was observed with sclerotia (74.29%), followed by mycelial inoculum (68.50%), when averaged across all cultivars. The lowest overall mean damping-off was found in colonized barley inocula (54.29%). The highest mean damping-off was found in Crystal 101RR (55.01%), followed by BTS8500 (54.39%). The lowest mean damping-off was observed in Maribo MA 504 (44.37%) and BTS 80RR52 (44.38%) (Table 2). The percent stand for each inoculum source and cultivar was directly and inversely correlated with percent damping-off (i.e, if damping-off was 60%, percent stand was 40%).

The effects of inoculum forms on root rot ratings at 56 dpi were significant (p<0.05)(Kruskal-Wallis chi-squared = 76.598, df = 3, p-value < 2.2e-16) (Table 4). Our analyzed p value was smaller that suggest there were significant difference among the inocula forms. It further suggests to run pairwise.wilcox.test in R. This helps to differentiate individual treatment or inocula forms. Mycelia and sclerotia were statistically non-significant. It showed p-value of 0.34 which was higher than 0.05. Sclerotia and colonized barley grains were statistically significant, since the pairwise p-value (0.0009) was smaller than 0.05 (Table 4).

**Table 4.** Comparison between the inocula forms through pairwise.wilcox.test (p<0.05).

| Pairwise inocula forms<br>and its p-value | Colonized barley grains | Autoclaved barley grains | Mycelia |
| --- | --- | --- | --- |
| Autoclaved barley grains | 1.20e-11 | - | - |
| Mycelia | 0.00017 | 1.50e-11 | - |
| Sclerotia | 0.0009 | 2.30e-11 | 0.34003<sup>NS</sup> |

NS, non-significant

**Table 5.**
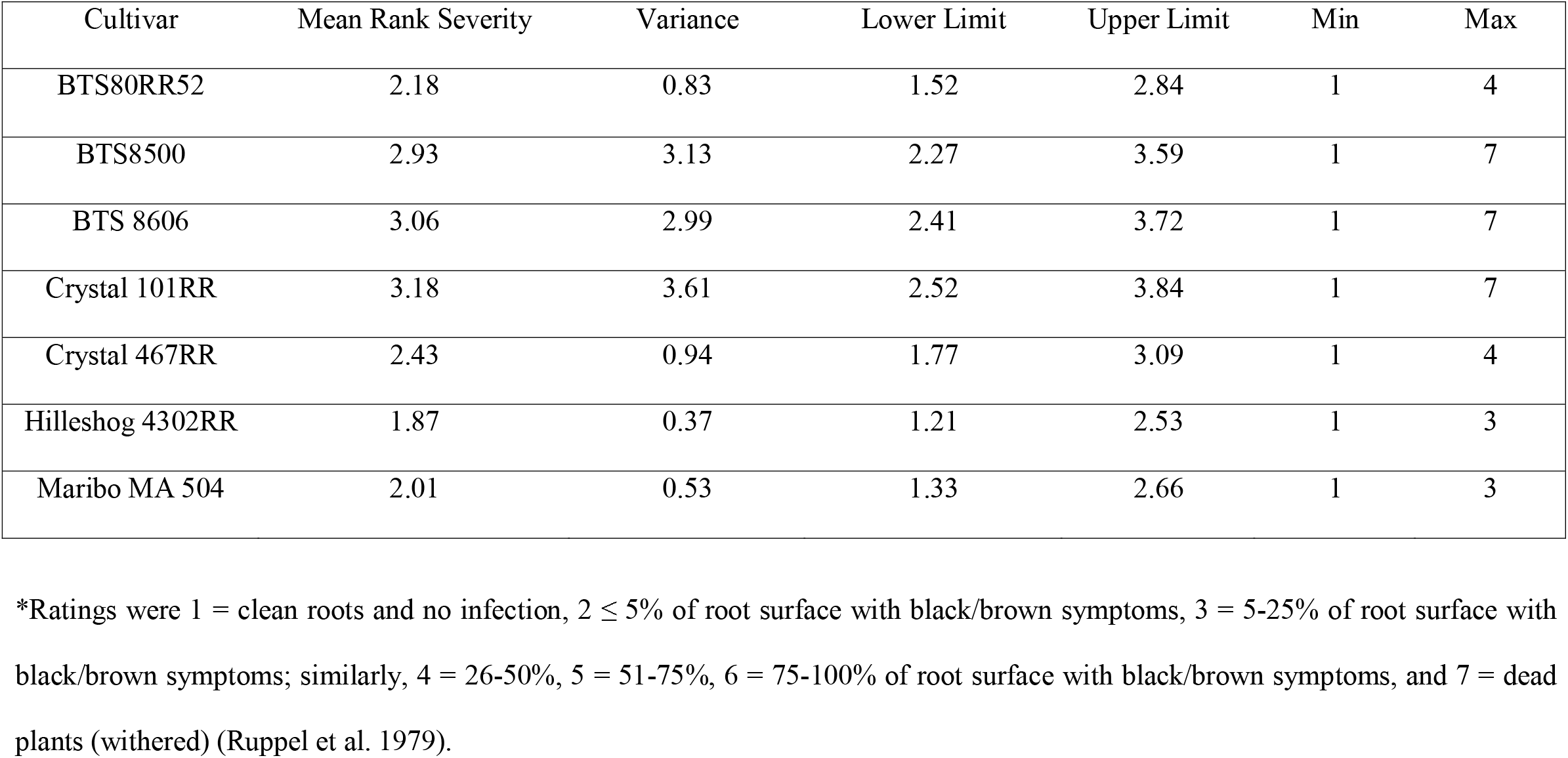
Non-parametric analysis for root rot ratings* at 56 dpi caused in cultivars

## 4. Discussion

Six culture media were evaluated to determine the most effective in vitro method of large-scale formations of sclerotia by *R. solani*. Hardar et al. (1981) demonstrated that sclerotia formation of *Sclerotium rolfsii* is induced in agar media within 3 days following scratching with scalpel to aerial mycelia. We demonstrated that ACV8 is a promising medium to produce *Rhizoctonia* sclerotia on a large scale, without the necessity of scratching the mycelia. The size of the sclerotia in our study varied significantly among the different culture media but the merits of inoculation potentials of different size of sclerotia remained similar (Garrett, 1956). This experiment showed an in vitro technique to prepare large scale sclerotia for pathogenic investigations.

Overwintering sclerotia, as well as melanized and moniloid mycelia are the primary source of infection during the seed germination stage in the field (Boland et al., 2004; Lee and Rush 1983). However, most previous pathogenetic studies were on adult beet plants using artificial inocula, mostly consisting of blended *Rhizoctonia* mycelia or *Rhizoctonia*-colonized cereal grains. Thus, those screening methods discounted the seed germination/seedling stage, which is the most vulnerable to stand losses due to damping off by *R. solani*. Our research also demonstrated that most commercial resistant sugar beet cultivars are highly susceptible to Rhizoctonia damping off at the seed germination stage.

In vitro inoculation study with three different types of *R. solani* inocula showed that the pathogenesis of sclerotia and the mycelial plug was better than colonized-barley grains in causing damping-off. This result demonstrated a novel approach of in vitro inoculation with three different forms of *R. solani* inocula in PDA media for studying host-pathogen interaction. This study demonstrated the use of sclerotia or mycelial plugs as a substitute for colonized barley/wheat/oat grains for evaluating the disease ratings of the commercial cultivars, as well as to simulate natural infection in the greenhouse.

In vivo inoculation studies in the greenhouse showed that all three forms of *Rhizoctonia* inocula were virulent and capable of damaging sugar beet plants. Inoculation with sclerotia showed more severe damping-off and root rot in the tested cultivars compared with the colonized-barley grains. This finding is in line with observations that sclerotia cause seed rot or pre-emergence damping-off by other research groups (Gaskill 1968; Naito and Makino 1995). This study was the first attempt to evaluate varietal resistance against *Rhizoctonia* damping-off on commercial cultivars using sclerotia and mycelial plugs. The results of this study suggest that sclerotia and mycelial plugs can be used as natural inocula to substitute for colonized barley grains in evaluating varietal resistance prior to release as a commercial cultivar. Recently, Liu et al. (2019) observed that sugar beet cultivars were highly susceptible to *R. solani* prior to attaining the six- to eight-leaf stage (4-5 weeks) after planting, regardless of the assigned level of resistance. The important response indicators of sugar beet cultivars include damping-off, root rot severity index, and stand count. Maribo MA 504 and BTS 80RR52 showed significantly lower damping-off compared to all other cultivars. Both cultivars showed the highest stand count and lowest root rot. This finding suggested that both Maribo MA504 and BTS 80RR52 can be used as resistant cultivars. Likewise, Crystal 101RR and BTS8500 were most susceptible among cultivars to damping-off. However, cultivation of Crystal 101RR and BTS 8500 in areas with existence of *R. solani* can be possible if the seed is protected by using fungicides. For example, use of a recommended dose of azoxystrobin in-furrow or another labeled fungicidal treatment is advised during the early growth stage, regardless of varietal resistance.

Sclerotia mediated damping-off in greenhouse studies showed significant variation in response among the cultivars. This emphasized the need to screen cultivars in the early stage of growth with sclerotia/mycelial plug inoculation to get a more reliable resistance rating than seen with adult plant screenings. Although growers in North America and Europe commonly use effective fungicides such as sedaxane or seeds coated with fungicides to control post-emergence damping-off, the seed companies need to consider age-dependent inoculation of plants for better evaluation of cultivars against susceptibility to *R. solani*.

In conclusion, an understanding of *Rhizoctonia* aggressiveness with natural inocula (sclerotia or mycelia) and screening of cultivars at the seed germination stage are essential for successful *R. solani* management. Future studies should evaluate interactions of cultivars at the seedling stage with other anastomosis groups of *R. solani* in order to minimize both stand and yield losses in sugar beets.

## Acknowledgments

Authors are very much thankful to D. Lakshman (USDA, ARS, Baltimore, Beltsville, MD 207052-350), Aiming Qi (School of Life and Medical Sciences, University of Hertfordshire, Hatfield, AL10 9AB, UK) and M. F. R. Khan (Department of Plant Pathology, North Dakota State University, Fargo, ND 58102, University of Minnesota, St. Paul, MN USA) for their suggestions, funding from USDA, and technical support.

